# Bridging the resolution gap in cryo-CLEM by introducing cryo-SXT: cryo-CLXEM

**DOI:** 10.1101/2025.09.05.673626

**Authors:** Johannes Groen, Anastasia Gazi, Sergey Kapishnikov, Anne Brelot, Matthijn Vos, Jost Enninga, Eva Pereiro, Anna Sartori-Rupp

## Abstract

Cryo-imaging in cellular biology provides the means to visualize the cellular interieur at close-to-native conditions. A cornerstone in the field has been cryo-correlative light and electron microscopy (cryo-CLEM), with cryo-visible light fluorescent microscopy (cryo-VLFM) providing the specificity by tagging macromolecules or structures and cryo-electron tomography (cryo-ET) for the structural details at molecular level. The large resolution gap between these techniques, however, is limiting this correlative workflow as cryo-ET targets are often smaller than the resolution limit of cryo-VLFM. Here we introduce cryo-soft X-ray tomography (cryo-SXT) as an intermediate step that can compensate for the partial view and limited resolution of cryo-VLFM by providing invaluable cellular context information in 3D to the cryo-ET dataset within an integrated workflow.

## Main text

Correlative multimodal imaging is a powerful tool for the study of complex systems, such as biological ones^1–5^. Correlating data obtained on the same sample by different modalities gives complementary information and supporting views, significantly promoting a more comprehensive interpretation of the studied cellular process. Major challenges for correlative multimodal imaging have been the compatibility of sample preparation and support, the drastic increase in complexity as each stage adds damage risk that can hinder subsequent steps, and the need to keep the workflow within a radiation dose limit, depending on the required resolution.

In cellular biology, correlative light and electron microscopy (CLEM) has been performed for many years at room temperature (RT)^6^. Recently, with the emergence of direct electron detectors and advanced image processing algorithms, cryo-ET has gained momentum and thus cryo-CLEM has set off in many laboratories^7,8^. It is now possible to generate 3D cryo-ET datasets at molecular resolution in near-native conditions. However, to generate high resolution (*i*.*e*. sub-nanometer) data, sample thickness (< 200 nm) and radiation dose (< 150 e/A^2^ ∼ 550 MGy^9,10^) need to be controlled^11^. Thus, for most unicellular or complex biological samples thinning is required, nowadays mostly done by cryo-focused Ion Beam (FIB) milling^12^. Cryo-VLFM is employed for targeting sites of interest, although inefficiently due to the resolution limitation and insufficient spatial accuracy for precise localization of the fluorescently labelled structures of interest, especially in the axial dimension. When aiming for small (<200 nm) structures, cryo-lamellae are often milled at the wrong depth, missing the target partially or completely.

Here we introduce cryo-SXT^13,14^ as an intermediate step, which can provide precious information regarding the 3D cellular context of up to 10 µm thick samples otherwise lost during cryo-FIB-milling.

With both lateral and axial resolution being higher than any existing cryo-VLFM modality, cryo-SXT provides the accuracy needed to optimize lamellae milling^15^. In cryo-SXT at the so-called “water window” energy region (284–543 eV), oxygen-based structures absorb little, while carbon-based structures absorb strongly the incoming photons. This provides a natural contrast of the cellular ultrastructure without additional staining. Moreover, intact cells can be imaged as a whole, without the need for sectioning. Cryo-SXT has been used in a correlative mode with cryo-VFLM (correlated light and X-ray microscopy, cryo-CLXM)^4,16–22^, and more recently in multimodal experimental setups also alongside cryo-CLEM^5,23^ or others^3^. Utilizing the three modalities together on the same cell within an integrated workflows (cryo correlative light, X-ray and electron microscopy: cryo-CLXEM), however, has to our knowledge never been reported yet.

Sample preparation is highly compatible for the three modalities as they use the same support and cryo-fixation approach. However, as radiation damage is induced by both X-rays^24,25^ and electrons^26,27^, their combination requires special care and evaluation of the resolution-dose tradeoff for imaging the sample. In cryo-soft X-ray imaging at Mistral (ALBA)^28^, a photodiode is used to measure the photon flux reaching the sample at a given energy and the percentage of photons absorbed is determined by the difference between the transmitted signal with and without the sample. In EM, what is commonly referred to as the electron dose describes the electron fluence - the number of electrons per unit surface to which the sample is exposed (expressed in e^−^/Å^2^). This value can be translated into an equivalent radiation dose, via a voltage-dependent conversion factor (see Table 2, online method) to be represented in MGy^10^. While there are various types of interactions^29,30^, soft X-rays mainly interact through photoelectric absorption, although a small portion is elastically scattered (Rayleigh Scattering) while electrons mainly interact through inelastic (Compton) scattering. In both cases, the absorbed dose depends on the composition and thickness of the sample, among other parameters related to the incoming beam (see online method). Despite the different interaction, the visible radiation effects are similar^31^, and a previous exposure can hasten the onset thereof. Hence, it can be assumed that a sample has a certain “dose-budget” that can be spent before visible damage to the ultrastructure becomes evident, depending on the desired resolution, and well before sublimation in the form of bubbling due to radiolysis of water molecules occurs^32^. Notably, the lack of visible damage does not imply its absences; molecular bonds will be broken at much earlier timepoints^33,34^. In our calculations, considering a goal of 5 nm resolution or better with cryo-ET^35^, we determined a total dose for cryo-SXT of a few tens of MGy^25,36^, corresponding approximately to 10-15% of the maximum dose-budget for vitrified ice sublimation. This was then considered for the subsequent cryo-ET acquisition (see online method).

**Table 1.** Parameters definition

| X-ray Microscopy | Definition/units | Electron Microscopy (EM) | Definition/units |
| --- | --- | --- | --- |
| **Radiation Dose** | $D[\text{MG}_y] = \text{Amount of energy deposited on the sample per unit mass}$ | **Radiation Dose** | $D = \text{Incident electron charge per unit area}$ |
| **Energy of the source** | $E(\text{eV})$ | **Energy of the source** | $E(\text{keV})$ |
| **Flux** | $n^\circ \text{ ph/s}$ | **Flux** | $n^\circ \text{ electrons/s}$ |
| **Flat field** | $f [\text{counts/s}]$ | **Camera gain reference** | Counts |
| **Flux/unit area** | $N \left[ \frac{\text{ph}}{\mu\text{m}^2 \text{ s}} \right]$ | **Flux/unit area (Dose rate)** | $N = \left[ \frac{e^-}{px^2 \text{ s}} \right]$ |
| **Fluence**<br>**= flux/unit area · t**<br>**= N · t** | $F \left[ \frac{\text{ph}}{\mu\text{m}^2} \right]$ | **Fluence (Exposure Dose)**<br>**= flux/unit area · t**<br>**= N · t** | $F = \left[ \frac{e^-}{\text{\AA}^2} \right]$ |

**Table 2.**
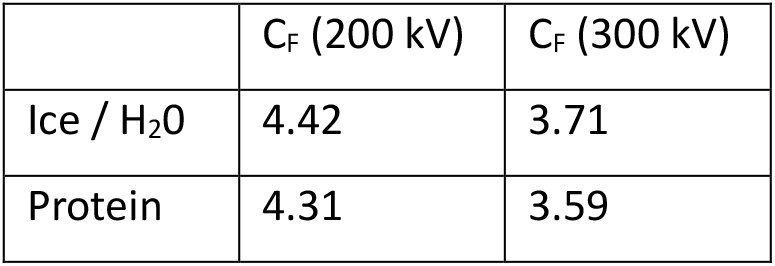
voltage-dependent conversion factor (C_F_) ^11^

**Table 3:** DOSE SUMMARY

|  | Soft-X rays |  | EM |  |
| --- | --- | --- | --- | --- |
| Pixel size | 13 nm |  | 3.104 Å |  |
| Exp time | 1 s |  | 1.77 s |  |
| Before sample | N<sup>b</sup> | 6.7 · 10<sup>7</sup> ph/ μm<sup>2</sup>/s | N<sup>b</sup> | 25 e<sup>-</sup>/pix<sup>2</sup>/s |
|  | F<sup>b</sup> | 6.7 · 10<sup>7</sup> ph/ μm<sup>2</sup> | F<sup>b</sup> | 4.78 e<sup>-</sup>/Å<sup>2</sup> |
|  | D<sup>b</sup><sub>TOT</sub> | 20.52 e<sup>-</sup>/Å<sup>2</sup> | F<sup>b</sup><sub>TOT</sub> | 110 e<sup>-</sup>/Å<sup>2</sup> |
|  |  | 76.15 MG<sub>y</sub> |  | 407 MG<sub>y</sub> |
|  | N<sup>a</sup> | 3,7 · 10<sup>7</sup> ph/ μm<sup>2</sup> /s | N<sup>a</sup> | 4.87 e<sup>-</sup>/pix<sup>2</sup>/s |
| After<br>sample | F<sup>a</sup> | $3,7 \cdot 10^7 \text{ ph/}\mu\text{m}^2$ | F<sup>a</sup> | $0.89 \text{ e}^-/\text{\AA}^2$ |
| | D<sup>a</sup><sub>TOT</sub> | $9.23 \text{ e}^-/\text{\AA}^2$ | F<sup>a</sup><sub>TOT</sub> | $20.70 \text{ e}^-/\text{\AA}^2$ |
|  |  | 34.23 MG<sub>y</sub> |  | 76.26 MG<sub>y</sub> |
|  | D<sub>TOT</sub> | 41.92 MG<sub>y</sub> | F<sub>TOT</sub> | 330.74 MG<sub>y</sub> |
| | Absorbed | $11.30 \text{ e}^-/\text{\AA}^2$ | Absorbed | $89.37 \text{ e}^-/\text{\AA}^2$ |

To briefly summarize the experimental procedure (Figure 1, supplementary data 1), adherent cells were grown on gold EM grids and plunge-frozen. Fluorescence labeling of the nucleus, mitochondria and acidic compartments was performed shortly before plunging. Vitrified samples were then imaged in a cryo-confocal microscope to, firstly, make a 2D map of the grid to assess the quality in terms of grid damage, ice quality and cell density and secondly, to identify and image suitable cells. Samples were then sent to the cryo-SXT Mistral beamline (ALBA Synchrotron, Spain) and loaded into the microscope. With their in-line visible light microscope (VLM) a 2D map is acquired and correlated with the confocal map to identify previously imaged cells. A single cryo-SXT tomogram per cell was acquired of up to 10 cells per grid, keeping tight control on the dose reducing the photon flux when required. It is important to note that balance must be found between exposure and resulting contrast (i.e. SNR) to avoid losing the contextual information provided by cryo-SXT. Post-acquisition, samples were shipped back to Institut Pasteur and loaded into the cryo-focused ion beam scanning electron microscope (FIB-SEM). Imaged cells were identified, and ∼300 nm thin lamellae were cut, aiming for the area previously imaged by cryo-SXT. The samples were then loaded into a cryo-transmission electron microscope (cryo-TEM) and tilt series of features of interest were acquired (see online method and supplementary data 2 for more details).

**Figure 1.**
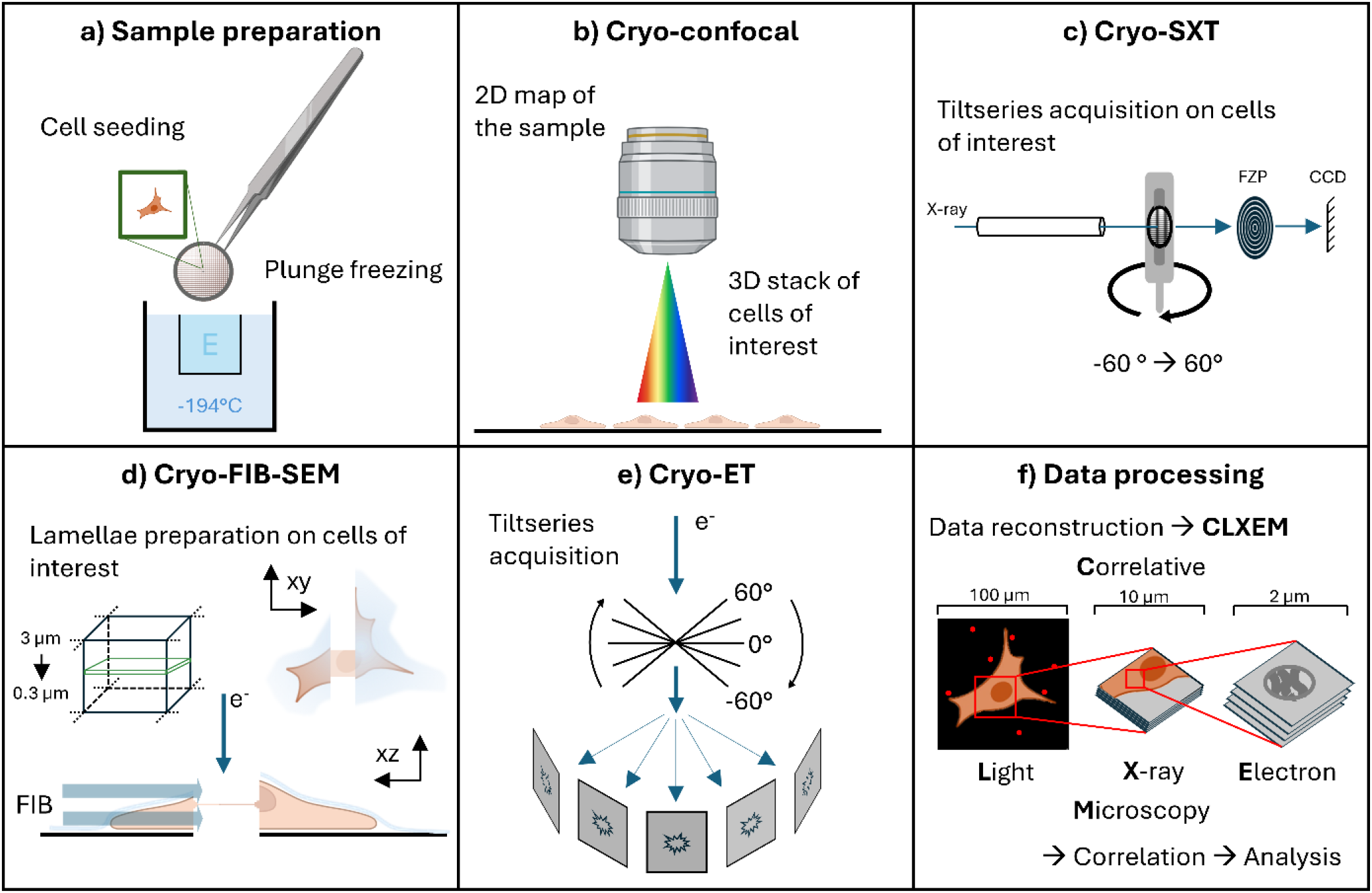
Complete cryo-CLXEM workflow. Cells are seeded on the grids and plunge frozen in liquid ethan (E, a). The samples are imaged by cryo-confocal microscopy (b), cells of interest are identified and z-stacks are acquired. Samples are then shipped to the synchrotron to be loaded in dedicated sample holders and cryo-SXT unidirectional tilt series are acquired (c). Samples are then loaded into a cryo-FIB-SEM, in which a focused ion beam (FIB) is used to trim away from the previously imaged cells of interest (d). Using a cryo-TEM, bidirectional tiltseries are acquired in the prepared lamellae (e). Finally, all the data are reconstructed and correlated before further analysis (f).

Figure 2 shows the full correlation from the 3 modalities in the same area. The fluorescent 1 µm microspheres can easily be identified in the cryo-confocal and cryo-SXT datasets. Using this information, the area of the cryo-SXT tomogram (in X and Y) could be localized within the cryo-FIB-SEM. At the cryo-TEM, medium magnification maps of the lamellae were acquired and sites for the acquisition of tilt series were chosen based on distinct features, like the two shown examples: mitochondria and multivesicular bodies, which can be easily identified in the cryo-SXT tomograms, were also labelled by fluorescent markers. Precise correlations were performed afterwards. An estimate of the achieved (half-pitch) resolution was made by Fourier shell correlation even/odd (cut-off at 0.25^35^) and were found to be ∼55 and ∼70 nm for the cryo-SXT volumes (with 25 and 40 nm Fresnel Zone Plates lenses, respectively), and ∼5 nm for the cryo-ET volume (see supplementary data 3). In addition to the tilt series, we acquired still-images to measure the allowed cumulative dose on the target. For this we took a series of 70 images at 0° tilt, resulting in a total (exposure) dose of up to ∼200 e/A^2^ (∼740 MGy) on the sample (see supplementary data 4), which did not show any apparent damage to visible membrane structures.

**Figure 2.**
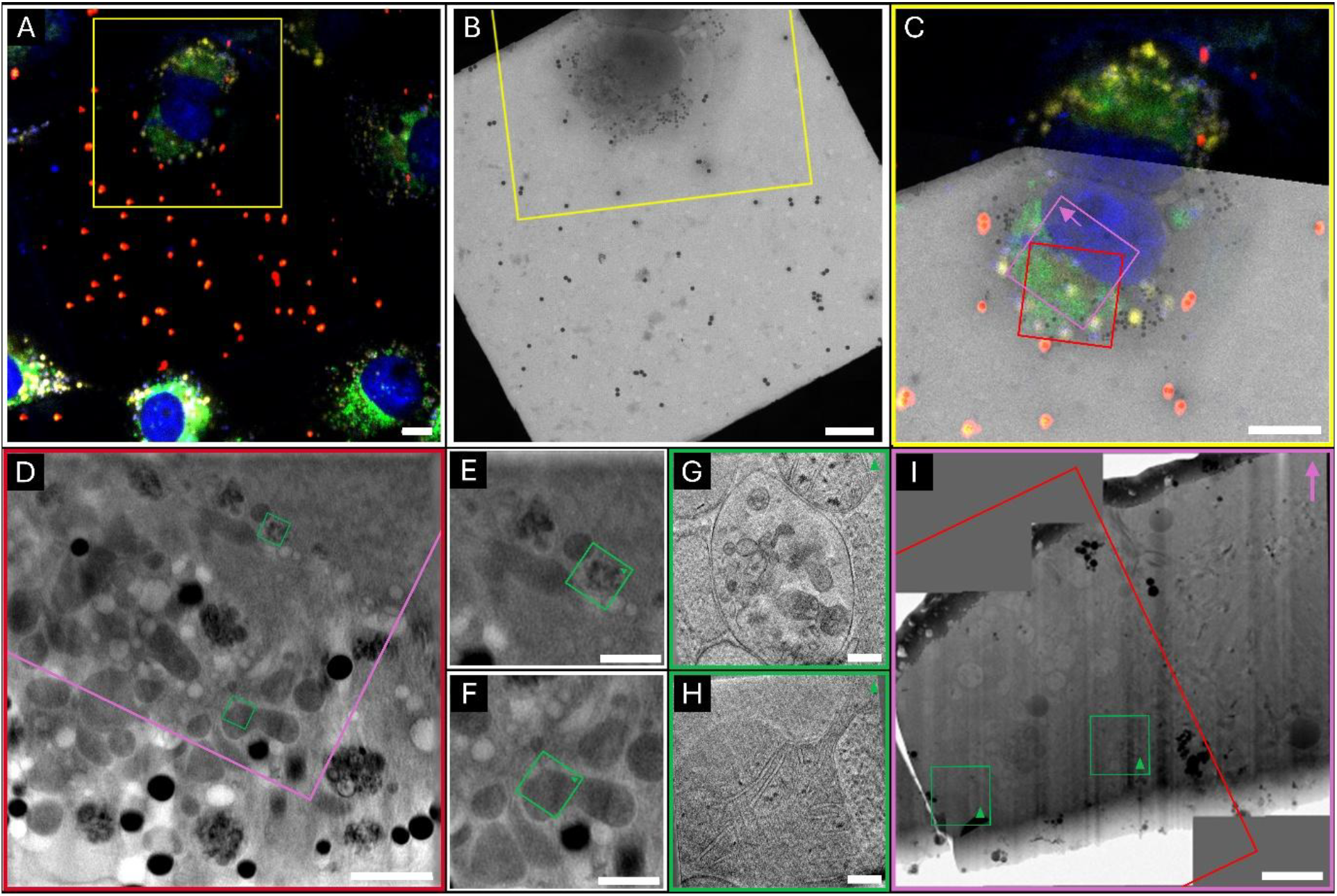
Results of the entire cryo-CLXEM pipeline. A shows the Z-projection of the confocal stack acquired at the cryo-Stellaris 8 confocal microscope. Blue – Hoechst, Green – Mitotracker Green, Red – Fluorescent Microspheres (1 µm), Yellow –Lysotracker deep red. B shows the 2D cryo-SXT mosaic of the same grid square. The yellow square in A and B are shown overlayed in C. The red square corresponds to the cryo-SXT tomogram area of which a single slice is shown in panel D (see supplementary video 1). The pink square corresponds to the lamella position, with the arrow indicating the direction (panel I). The green squares highlight the area of the cryo-ET acquisitions. In E and F single slices from the reconstructed cryo-SXT tomogram, showing a multivesicular body (see supplementary video 2) and part of a mitochondrion, respectively, are shown, which can also be observed by cryo-ET in panels G and H. Scalebars: A, B, C: 10 µm; D: 2 µm; E, F: 1 µm; G, H: 200 nm; I: 1 µm

Herein, we have developed an integrated cryo-CLXEM workflow demonstrating that acquisition of cryo-electron tomograms for the visualization of eukaryotic cells is possible after cryo-confocal and cryo-SXT imaging. Although the throughput remains limited in our study - with approximately 15% of the cryo-SXT imaged samples targeted by cryo-ET, we believe that the potential of this cryo-correlative multiscale workflow is evident.

The estimated resolutions are in an acceptable range considering the reduced dose and SNR used in this proof-of-concept study. For studying rare events which are inherently difficult to locate, cryo-CLXEM will allow precise targeting, in addition to securing the cellular context for cryo-ET data interpretation. Further investigations using sub-tomogram averaging would need to be performed on known macromolecules (e.g. *in situ* ribosomes) to evaluate the ultimate resolution achievable. Note that one of the major bottlenecks is the targeting at the cryo-FIB-SEM system, which directly correlates to the throughput. The small field of view (FoV) contained few to no correlative markers, which in addition were mostly indistinguishable from the accumulated ice-crystal contamination from the FIB-angle point of view. With future technological developments, we expect this to improve. Meanwhile, intermediate solutions, like using more distinguishable correlative fiducial markers and/or using correlation software like 3DCT^37^ could enhance the workflow. Moreover, we suggest the development of software tools to allow calculating in advance the total dose a specific sample could stand for a given spatial resolution, following what other scientific communities have done, *e*.*g*. the protein crystallography field^38^.

Overall, the addition of cryo-SXT provides a wealth of complementary information to classical cryo-ET datasets. The larger FoV and depth, that allow visualizing the local cellular environment, can be crucial for the interpretation of structures within the lamellae. Additionally, with scientists experiencing the need to go to more complex models, like tissue sections or organoids, cryo X-ray imaging can provide a real benefit over cryo-VLM when it comes to targeting areas of interest.

## Acknowledgements

Cryo-SXT data were collected at Mistral beamline at Alba Synchrotron during experimental sessions 2023087704. This work was supported by grants from the Programme Fédérateur de Recherche 6 (PFR6) SARS-CoV-2 & COVID-19. We acknowledge the cryo-ET expertise and assistance of the Institut Pasteur’s NanoImaging Core facility, created and supported by a PIA grant (EquipEx CACSICE: ANR-11-EQPX-0008). The DIHP unit is part of the LabEx IBEID and of the LabEX MI. Furthermore, JE acknowledges funding by the ANR PRC RabReprogram.

## Contribution

J.G. performed all the experimental steps and wrote the manuscript. A.S.R., E.P. and J.G. conceived the study. A.S.R. supervised the study. E.P. collected the cryo-SXT data at Mistral (ALBA). A.S.R and E.P. provided the dose calculations and scientific discussions. J.E. provided edits to the paper and financial support. A.G and S.K. provided scientific discussions. A.B. provided financial support and M.V. provided facility support.

### Online method

#### Sample preparation

VeroE6 cells were maintained in DMEM GlutaMax medium supplemented with 10% FBS and 1% Pen/Strep. 150.000 cells were seeded in a 35 mm ibidi culture dish, on 200 Mesh 1/4 holey carbon gold EM-grids from Quantifoil. Hoechst nuclear dye, Mitotracker green and Lysotracker deep-red (all from Thermo Fischer Scientific) were added shortly before plunge-freezing, considering the manufacturers recommended concentration and incubation time, washed with PBS, and returned to complete medium until freezing. The grids were frozen using the Leica EMGP plunge-freezer, keeping the humidity in the chamber above 70% and adding 3 µl of 1 µm fluorescent microsphere (red FluoSpheres™, Invitrogen) to the grids before blotting. Grids were blotted for 5 seconds, plunged into liquid ethane kept at -180°C, clipped into the specialized Thermo Fischer autogrids for cryo-FIB-milling, and stored in liquid nitrogen until further use.

#### Cryo-confocal imaging

Samples were loaded into the cryo-Stellaris 8 confocal system (Leica Microsystems) equipped with a HC PL APO 50x/0.90 cryo objective, a white light laser (WLL) and three HyD detectors. A first 2D widefield map was created of the entire grid to assess the quality of the grid in terms of damage, number of cells in suitable positions and quality of the ice. Then cryo-confocal stacks were acquired of up to 10 cells of interest per grid, with the pinhole set to 1 Au, the zoom set to 1.7x to have the FOV be one grid square only and the z-step was determined using the system-optimization feature.

#### Cryo-soft X-ray tomography (cryo-SXT) imaging

Samples were shipped to Mistral Beamline at the Alba Synchrotron in Spain using a dry shipper (CX100). There they were loaded into their dedicated dual tilt sample holders ^1^ and inserted into the cryo-SXT microscope ^2^. The magnification was chosen to give an effective pixel size of 13 nm for the 40 nm Fresnel Zone Plate (FZP) and 11 nm for the 25 nm FZP. First, the photodiode was inserted to measure the real flux at the sample position. Then, the sample was loaded and a brightfield and fluorescence map were acquired using the in-build visible light microscope. Then, using the 2D fluorescence map from the cryo-confocal system, the cells that were previously imaged were identified. Only one X-rays tomogram per cell was acquired, as the X-ray radiation exposure extends beyond the FOV. The dose on the sample was controlled by reducing the gap of the entry and exit slit of the monochromator. X-ray images were acquired every 1 degree, up to +/-65 degrees, if possible, for a total of up to 131 images. The samples were then unloaded and shipped back to Institut Pasteur Paris, France.

#### Cryo-lamellae preparation

For the cryo-Focused Ion Beam (FIB) milling, samples were loaded into the Thermo Fischer Aquilos 2 system. Using the Maps-software, the acquired SEM overview map of the grid was aligned with the previously acquired visible light microscopy maps to identify the areas imaged by cryo-confocal and cryo-SXT. Using the 2D-X-ray mosaic images, the exact area within the grid square was identified and marked. Before starting the milling process an organometallic platinum layer was deposited on the grids for 75 seconds using the gas injection system. The Auto-TEM software was used for automatic lamellae cutting, from 1 nA for initial rough milling, going down gradually until 30 pA for the final fine polishing step, aiming for a final thickness of 300 nm. After completion, the lamellae were inspected manually and polished further if any curtaining was visible.

#### Cryo-EM

Samples were loaded into the 300 kV cryo-TEM Titan Krios, equipped with a Falcon 4i direct electron detector (Thermo Fisher Scientific) and a Selectris X energy filter (Thermo Fisher Scientific), using the autoloader system. Datasets were collected using SerialEM (Version 4.1.4) ^3^ with the energy filter at 10 eV (zero loss), objective aperture of 100 μm and a pixel size of 3.104 Å. The exposure time was chosen to have a maximum of 110 e/Å^2^ for the full tilt series. The tilt series were acquired utilizing a dose-symmetric scheme, acquiring an image every 3°, from 70° to -50°, starting at 10°, for a total of up to 41 images at defocus set to -2 μm.

#### Data processing

Cryo-ET frames were aligned using MotionCor 2 within the Scipion package ^4^. Both cryo-SXT and cryo-ET tilt series were aligned with AreTomo ^5^ and reconstructed using either the SIRT algorithm of Tomo3D ^6^ for the cryo-SXT datasets and using the Filtered Back Projection with the SIRT-like filter of IMOD ^7^ for the cryo-ET datasets. Initial correlation of the data was done with scNodes ^8^ and for a finer correlation we utilized the ecCLEM plugin ^9^ in ICY ^10^.

#### Resolution estimation

The resolution was estimated by Fourier shell correlation (FCS) of the split tilt series, as reported previously (Carrascosa et al., 2009). For the cryo-SXT datasets, as no frames are collected, the tilt series were divided into even and odd tilts. For the cryo-EM datasets, the frames were divided. AreTomo was used for the alignment and the datasets were reconstructed using the WBP algorithm with the thickness limited to include only cellular features. For the calculation the resolution FSO protocol of XMIPP3 was used withing the Scipion environment and the Global FSC output was used.

#### Doses estimation

where:

- e^-^ = electron
- 1 eV = 1,6 · 10^-19^ J = energy of 1 e^-^ under a potential difference of 1V
- px = pixel size

### X-rays radiation-dose formula

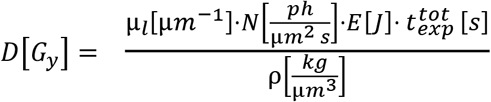

where:

- D = radiation dose measured in Grey = G_y_[J/kg], which is the amount of energy deposited through inelastic scattering per unit mass of specimen, measured in Joule per kilogram
- ρ is the density of the material, where ρ ∼ 0.94 g/cm^3^ is the average value for amorphous ice
- μ_*>l*_ is the linear absorption coefficient, where µ_l_ ∼ 0.1 µm^-1^ is the average value for amorphous ice
- N = Flux/unit area = (number of photons/s)/unit area
- ph = photon
- E [J] = energy of the X-rays source measured in Joule
- t^tot^_>exp_ = total exposure time

### Electron radiation-dose formula

Conversion of e^-^/Å^2^ to *MG*_*>y*_: [*MG*_*>y*_] = C_F_ · [e^-^/Å^2^]

### Parameters for cryo-soft X-rays tomography half-dose acquisition at Mistral Beamline

- Source energy: E = 520 eV = 8.33 x 10-17 J
- Fresnel zone plates (FZP) = 40 nm
- Pixel size (px) = 13 nm
- Flux (total flux at 520 eV at sample position with ES= 30 µm and XS=15 µm) = 2.08 · 10^10^ ph/s **(measured with photodiode)**
- XS (for acquisitions) = 8 µm **→** Flux (for acquisitions) = 53% of Flux (at sample position) = 1.11 · 10^10^ ph/s
- Field of View (FoV) = 13 um x 13 um = 169 µm^2^
- Total n° projections = 131 t_exp_ = 1 s (Exposure time per projection)
- µ_l_ ∼ 0.1 µm^-1^
- ρ ∼ 0.94 g/cm^3^ = 0.94 · 10^-15^ kg/µm^3^

**Calculation of Fluence F = N · t [ph/µm^2^]** (equivalent to Exposure Dose E_d_ [é/A^2^] in EM)

#### Before the sample

Flat field: f^b^ = 39000 cts/s (average value on all tilt series projections)

Fluence: F^b^ = 6.7 · 10^7^ ph/s µm^2^ (fluence reaching the sample per FoV per projection)

Total Fluence: F^b^_TOT_ = 131 · F^b^ = 0.86 x 10^10^ ph/µm^2^ per tilt series

Total Dose: D^b^_TOT_ [MG_>y_]= µ_l_/ρ · F^b^_TOT_ · E · t_exp_ = **76.20 MG**_**>y**_

#### After the sample

Flat field: f^a^ = 17540 cts/s (average value on all tilt series projections)

Fluence: F^a^ = 3.7 · 10^7^ ph/µm^2^ (fluence reaching the sample per FoV per projection) Total

Fluence: F^a^_TOT_ = 131 · F^a^ = 0.48 · 10^10^ ph/µm^2^ per tilt series

**→** 55% adsorbed fluence, supposing linear response of the FZP.

**→** total adsorbed dose = 0.55 · D_TOT_ [MG_>y_]= **41.92 MG**_**>y**_

### Parameters for full-dose cryo-electron tomography acquisition on the Titan Krios

- Source energy: E = 300 keV
- Total n° projections = 41
- Pixel size (px) = 3.104 Å
- t_exp_ = 1.77 s (Exposure time per projection)
- N (Dose rate) = 25.58 e^-^/px^2^/s
- F (Fluence, Exposure dose) = 4.78 e^-^/Å^2^
- D_TOT_ (Total Radiation Dose per tilt series) = (F · 41) /t_exp_ = 110.48 e^-^/Å^2^

### Calculation of Fluence/Exposure Dose F[é/A^2^] for EM (equivalent to Fluence F[ph/µm^2^] for X-rays)

#### Before the sample

Fluence: F^b^ = 4.78178 e^-^/Å^2^ (fluence reaching the sample per FoV per projection)

Total Fluence: F^b^_TOT_ = 41 · F^b^ /t_exp_ = 110 e^-^/Å^2^ per tilt series

Total Dose: F^b^_TOT_ (e^-^/Å^2^) · 3.7 = **407 MG**_**>y**_

#### After the sample

Fluence: F^a^ = 0.89 e^-^/Å^2^ (fluence reaching the sample per FoV per projection)

Total Fluence: F^a^_TOT_ = 41 · F^a^ /t_exp_= 20.61 e^-^/Å^2^ per tilt series

Total Dose: F^a^_TOT_ (e^-^/Å^2^) · 3.7 = **76.26 MG**_**>y**_

**→** 81.3% adsorbed fluence

**→** Total adsorbed dose = 0.813 x D^b^_TOT_ [*MG*_*>y*_] = **330.74** MG_>y_ At 300 kV: [MG_>y_] = C_F_ · [e^-^/Å^2^], with C_F_ = 3.71, as biological material is mostly composed of H_2_0.

## Conclusions

To achieve resolutions better than 3 Å, exposure should be kept below 30 e^-^/Å^2 12^, or 111 MG_>y_. D^b^_TOT_ has to be kept to minimum to allow high resolution reconstructions and structure determination afterwards, keeping in mind that for a good cryo-SXT tomogram, 40-60% absorption provides good S/N ratio, without using denoising algorithms. Absorption depends highly on composition and thickness and photon flux will be different for each cryo-SXT source.

---

**Supplementary Figure 1.**
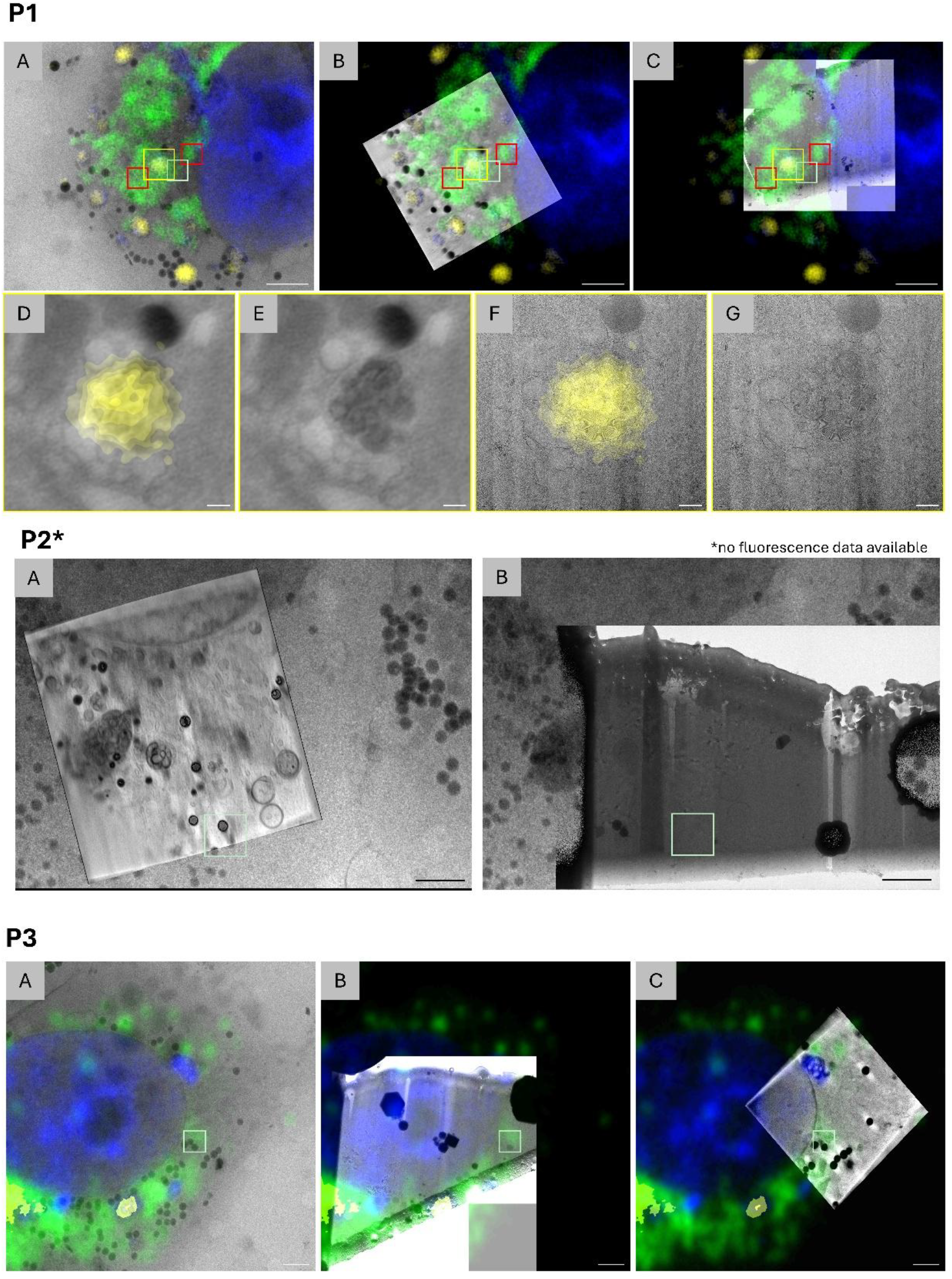
Additional areas with partial overlap of the cryo-SXT tomogram and a lamella. P1 corresponds to the correlation show in figure 1, highlighting the 3-modality-correlation that shows an acidic compartment (stained by lysotracker) that can be identified also in the cryo-SXT tomogram and the cryo-EM lamella. P2 and P3 show partially overlapping areas in which no tomograms were collected in the overlapping regions. Scale bars : P1-ABC, P2-AB, P3-ABC : 2 μm; P1-D-G : 200 nm

**Supplementary Figure 2.**
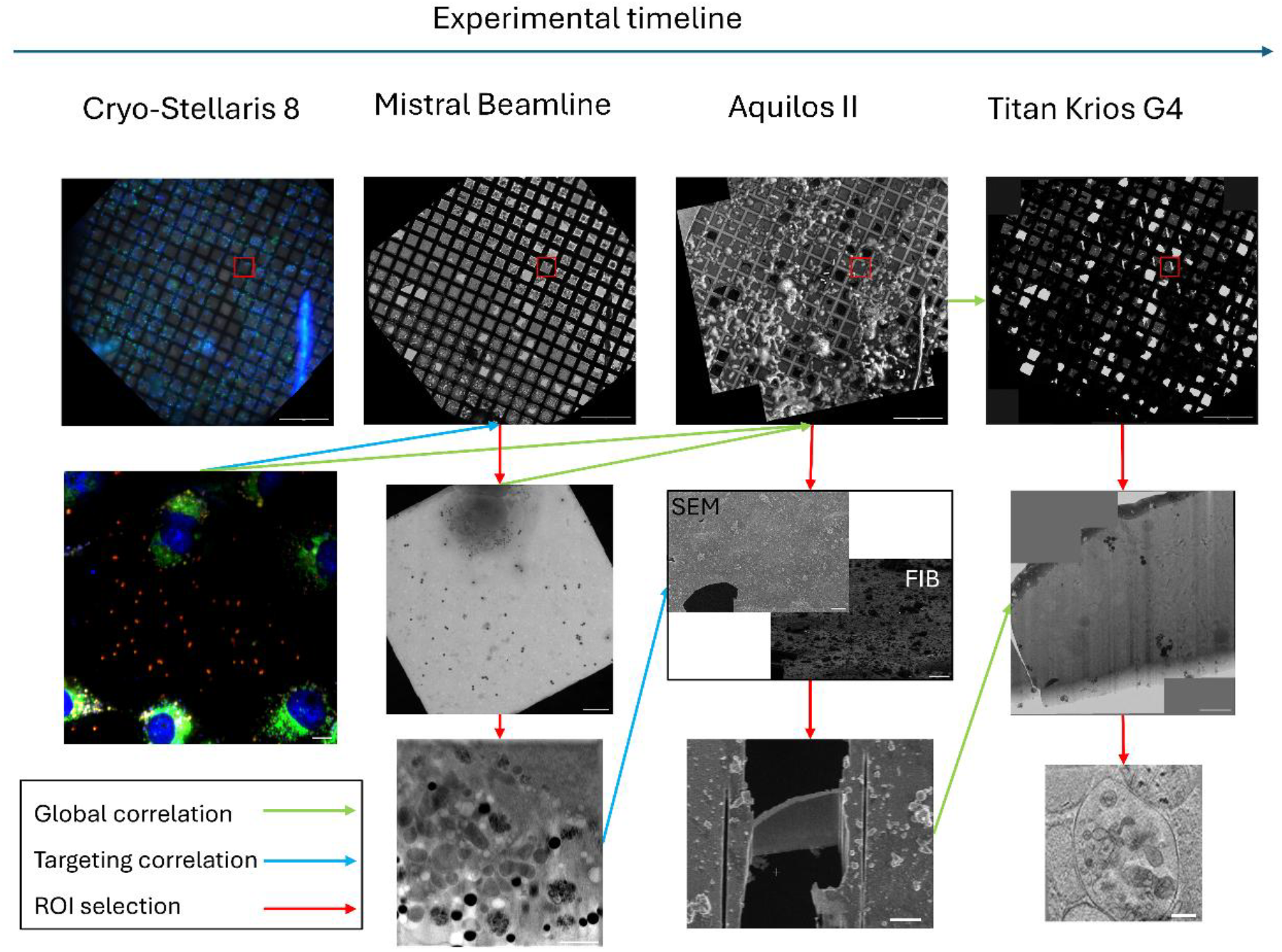
Timeline and correlations that have been performed. Green arrows highlight global correlations, used mostly for navigation purposes. Blue arrows show the targeting correlations that needed to be accurate for targeting. Red arrows show the manual ROI selection process, guided by the global correlations. Scale bars : top row : 50 μm; middle row, cryo-stellaris and Mistral : 10μm, Aquilos : 20 μm, Titan : 2 μm; bottom row, Mistral : 2 μm, Aquilos 5 μm, Titan : 200 nm

**Supplementary Figure 3.**
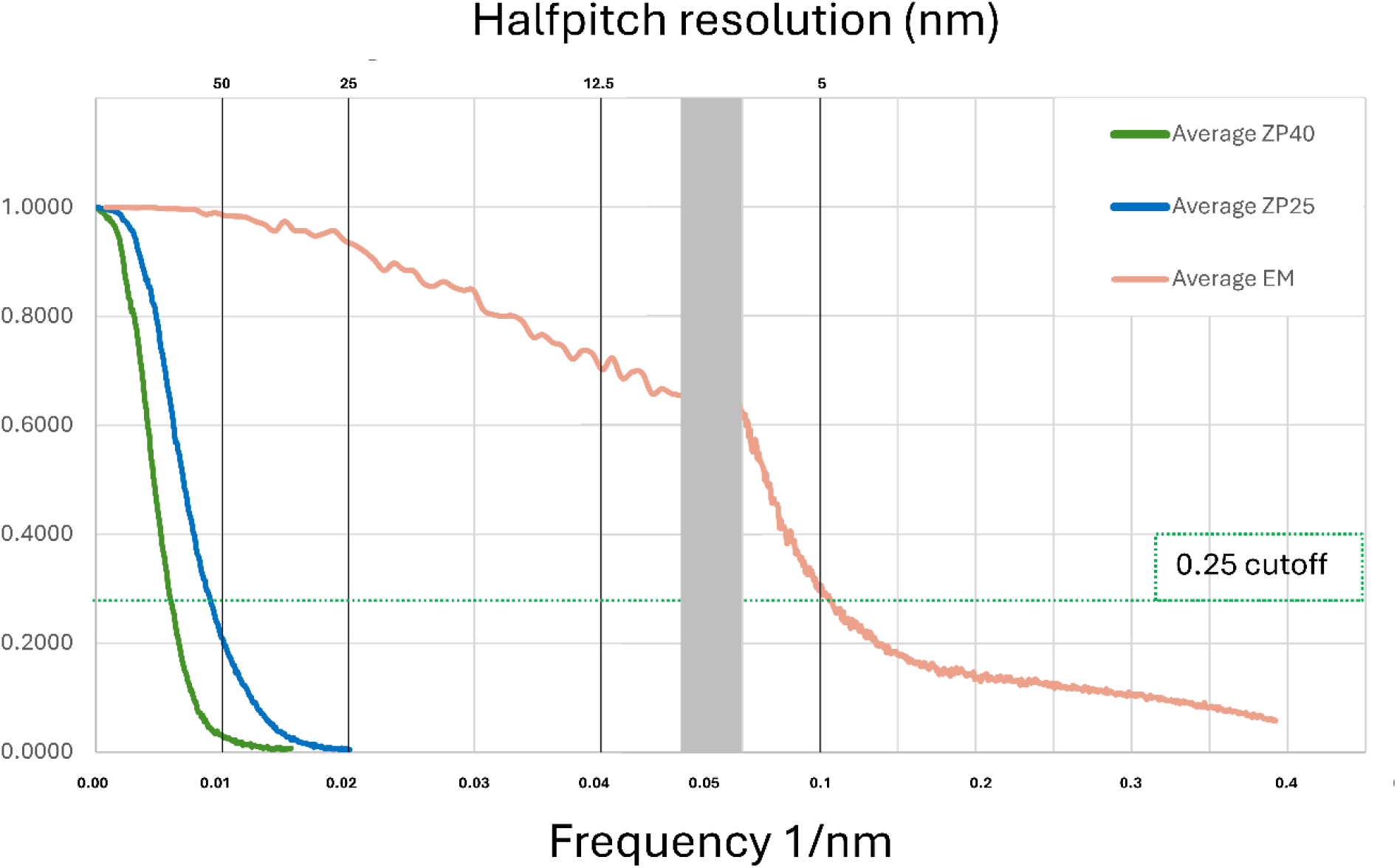
Averaged FSC curve showing the achieved resolution. FZP 40 nm : ∼80 nm; FZP 25 nm : ∼53 nm; EM: ∼5 nm Supplementary Figure 4 First and last images from a 70 image stack acquired from 3 areas that were previously imaged by cryo-SXT, highlighting the lack of visible radiation damage. Scalebars : 200 nm

**Supplementary Figure 4.**
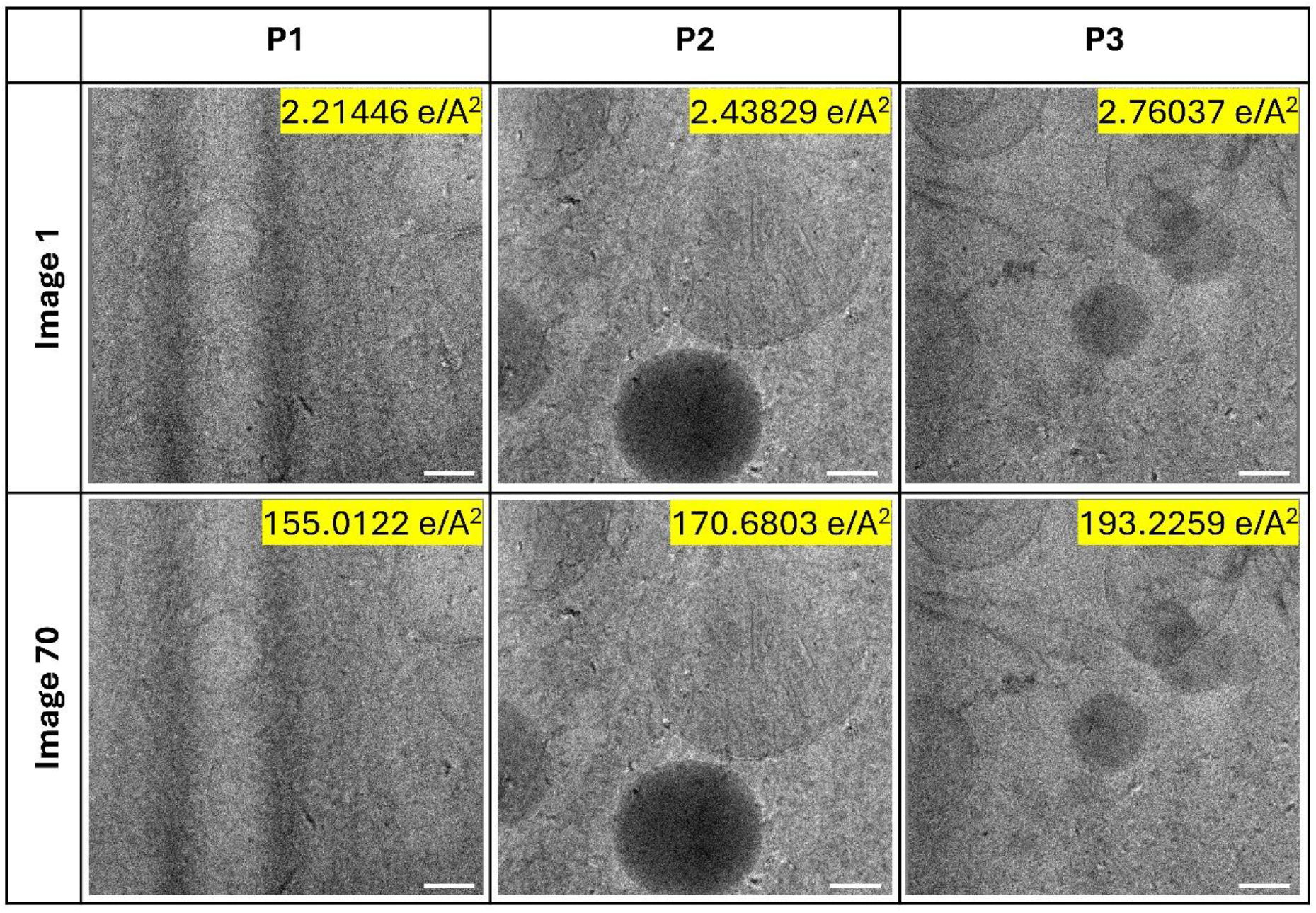
First and last images from a 70 image stack acquired from 3 areas that were previously imaged by cryo-SXT, highlighting the lack of visible radiation damage. Scalebars : 200 nm

## Notes

### Competing Interest Statement

Sergey Kapishnikov is part of SiriusXT, a company developing a cryo-SXT laboratory system. The other authors declare no competing interests.

## References

1. De Boer, P., Hoogenboom, J. P. & Giepmans, B. N. G. Correlated light and electron microscopy: Ultrastructure lights up! Nat Methods 12, (2015).

2. van Rijnsoever, C., Oorschot, V. & Klumperman, J. Correlative light-electron microscopy (CLEM) combining live-cell imaging and immunolabeling of ultrathin cryosections. Nat Methods 5, (2008).

3. Conesa, J. J. et al. Unambiguous Intracellular Localization and Quantification of a Potent Iridium Anticancer Compound by Correlative 3D Cryo X-Ray Imaging. Angewandte Chemie -International Edition 59, (2020).

4. Duke, E. M. H. et al. Imaging endosomes and autophagosomes in whole mammalian cells using correlative cryo-fluorescence and cryo-soft X-ray microscopy (cryo-CLXM). Ultramicroscopy 143, 77–87 (2014).

5. Mendonça, L. et al. Correlative multi-scale cryo-imaging unveils SARS-CoV-2 assembly and egress. Nat Commun 12, (2021).

6. Kukulski, W. et al. Correlated fluorescence and 3D electron microscopy with high sensitivity and spatial precision. Journal of Cell Biology 192, (2011).

7. Lučić, V. et al. Multiscale imaging of neurons grown in culture: From light microscopy to cryo-electron tomography. J Struct Biol 160, (2007).

8. Sartori, A. et al. Correlative microscopy: Bridging the gap between fluorescence light microscopy and cryo-electron tomography. J Struct Biol 160, (2007).

9. Henderson, R. Cryo-protection of protein crystals against radiation damage in electron and X-ray diffraction. Proc R Soc Lond B Biol Sci 241, 6–8 (1990).

10. Baker, L. A. & Rubinstein, J. L. Radiation damage in electron cryomicroscopy. in Methods in Enzymology vol. 481 (2010).

11. Hoffmann, P. C. et al. Structures of the eukaryotic ribosome and its translational states in situ. Nat Commun 13, (2022).

12. Wagner, F. R. et al. Preparing samples from whole cells using focused-ion-beam milling for cryo-electron tomography. Nat Protoc 15, (2020).

13. Schneider, G. et al. Three-dimensional cellular ultrastructure resolved by X-ray microscopy. Nat Methods 7, 985–987 (2010).

14. Parkinson, D. Y., McDermott, G., Etkin, L. D., Le Gros, M. A. & Larabell, C. A. Quantitative 3-D imaging of eukaryotic cells using soft X-ray tomography. J Struct Biol (2008) doi:10.1016/j.jsb.2008.02.003.

15. Schneider, G. Cryo X-ray microscopy with high spatial resolution in amplitude and phase contrast. Ultramicroscopy 75, (1998).

16. Carzaniga, R., Domart, M. C., Duke, E. & Collinson, L. M. Correlative Cryo-Fluorescence and Cryo-Soft X-Ray Tomography of Adherent Cells at European Synchrotrons. Methods Cell Biol (2014) doi:10.1016/B978-0-12-801075-4.00008-2.

17. Spink, M. C. et al. Correlation of Cryo Soft X-ray Tomography with Cryo Fluorescence Microscopy to Characterise Cellular Organelles at Beamline B24, Diamond Light Source. Microscopy and Microanalysis (2018) doi:10.1017/s1431927618014162.

18. Okolo, C. A. et al. Sample preparation strategies for efficient correlation of 3D SIM and soft X-ray tomography data at cryogenic temperatures. Nat Protoc 16, (2021).

19. Groen, J. et al. Correlative 3D cryo X-ray imaging reveals intracellular location and effect of designed antifibrotic protein-nanomaterial hybrids. Chem Sci 12, (2021).

20. Hagen, C. et al. Multimodal nanoparticles as alignment and correlation markers in fluorescence/soft X-ray cryo-microscopy/tomography of nucleoplasmic reticulum and apoptosis in mammalian cells. Ultramicroscopy 146, (2014).

21. Cinquin, B. P. et al. Putting molecules in their place. J Cell Biochem 115, (2014).

22. Hagen, C. et al. Correlative VIS-fluorescence and soft X-ray cryo-microscopy/tomography of adherent cells. J Struct Biol 177, 193–201 (2012).

23. Castro, V., Pérez-Berna, A. J., Calvo, G., Pereiro, E. & Gastaminza, P. Three-Dimensional Remodeling of SARS-CoV2-Infected Cells Revealed by Cryogenic Soft X-ray Tomography. ACS Nano 17, (2023).

24. Guo, L. et al. The study of radiation damage of yeast cells in Cryo-soft x-ray tomography. in Selected Papers of the Chinese Society for Optical Engineering Conferences held October and November 2016 (eds. Lv, Y., Le, J., Chen, H., Wang, J. & Shao, J.) 102551Q (2017). doi:10.1117/12.2268715.

25. Howells, M. R. et al. An assessment of the resolution limitation due to radiation-damage in X-ray diffraction microscopy. J Electron Spectros Relat Phenomena 170, (2009).

26. Egerton, R. F. Dose measurement in the TEM and STEM. Ultramicroscopy 229, (2021).

27. Karuppasamy, M., Karimi Nejadasl, F., Vulovic, M., Koster, A. J. & Ravelli, R. B. G. Radiation damage in single-particle cryo-electron microscopy: Effects of dose and dose rate. J Synchrotron Radiat 18, (2011).

28. Sorrentino, A. et al. MISTRAL: a transmission soft X-ray microscopy beamline for cryo nano-tomography of biological samples and magnetic domains imaging. J Synchrotron Radiat 22, 1112–1117 (2015).

29. Henke, B. L., Gullikson, E. M. & Davis, J. C. X-ray interactions: Photoabsorption, scattering, transmission, and reflection at E = 50-30, 000 eV, Z = 1-92. At Data Nucl Data Tables 54, (1993).

30. Reimer, L. & Kohl, H. Transmission Electron Microscopy Physics of Image Formation. Springer series in optical sciences vol. 51 (2008).

31. Baker, L. A. & Rubinstein, J. L. Radiation Damage in Electron Cryomicroscopy. in Methods in Enzymology vol. 481 371–388 (2010).

32. Leapman, R. D. & Sun, S. Cryo-electron energy loss spectroscopy: observations on vitrified hydrated specimens and radiation damage. Ultramicroscopy 59, (1995).

33. Henderson, R. The Potential and Limitations of Neutrons, Electrons and X-Rays for Atomic Resolution Microscopy of Unstained Biological Molecules. Q Rev Biophys 28, (1995).

34. Grant, T. & Grigorieff, N. Measuring the optimal exposure for single particle cryo-EM using a 2.6 Å reconstruction of rotavirus VP6. Elife 4, (2015).

35. Carrascosa, J. L. et al. Cryo-X-ray tomography of vaccinia virus membranes and inner compartments. J Struct Biol 168, 234–239 (2009).

36. Atakisi, H., Conger, L., Moreau, D. W. & Thorne, R. E. Resolution and dose dependence of radiation damage in biomolecular systems. IUCrJ 6, (2019).

37. Arnold, J. et al. Site-Specific Cryo-focused Ion Beam Sample Preparation Guided by 3D Correlative Microscopy. Biophys J 110, (2016).

38. Paithankar, K. S. & Garman, E. F. Know your dose: RADDOSE. Acta Crystallogr D Biol Crystallogr 66, (2010).

## References

1. Valcarcel, R. et al. New Holder for Dual-Axis Cryo Soft X-Ray Tomography of Cells at the Mistral Beamline. in Mechanical Engineering Design of Synchrotron Radiation Equipment and Instrumentation (2018).

2. Sorrentino, A. et al. MISTRAL: a transmission soft X-ray microscopy beamline for cryo nano-tomography of biological samples and magnetic domains imaging. J Synchrotron Radiat 22, 1112– 1117 (2015).

3. Mastronarde, D. N. Automated electron microscope tomography using robust prediction of specimen movements. J Struct Biol 152, (2005).

4. de la Rosa-Trevín, J. M. et al. Scipion: A software framework toward integration, reproducibility and validation in 3D electron microscopy. J Struct Biol 195, (2016).

5. Zheng, S. et al. AreTomo: An integrated software package for automated marker-free, motion-corrected cryo-electron tomographic alignment and reconstruction. J Struct Biol X 6, (2022).

6. Agulleiro, J. I. & Fernandez, J. J. Fast tomographic reconstruction on multicore computers. Bioinformatics 27, 582–583 (2011).

7. Kremer, J. R., Mastronarde, D. N. & McIntosh, J. R. Computer Visualization of Three-Dimensional Image Data Using IMOD. J Struct Biol 116, 71–76 (1996).

8. Last, M. G. F., Voortman, L. M. & Sharp, T. H. scNodes: a correlation and processing toolkit for super-resolution fluorescence and electron microscopy. Nature Methods vol. 20 (2023).

9. Paul-Gilloteaux, P. et al. eC-CLEM: flexible multidimensional registration software for correlative microscopies. Nat Methods 14, 102–103 (2017).

10. de Chaumont, F. et al. Icy: an open bioimage informatics platform for extended reproducible research. Nat Methods 9, 690–696 (2012).

11. Egerton, R. F. Dose measurement in the TEM and STEM. Ultramicroscopy 229, (2021).

12. Grant, T. & Grigorieff, N. Measuring the optimal exposure for single particle cryo-EM using a 2.6 Å reconstruction of rotavirus VP6. Elife 4, (2015).

